# Neural envelope tracking as a measure of speech understanding in cochlear implant users

**DOI:** 10.1101/469643

**Authors:** Eline Verschueren, Ben Somers, Tom Francart

**Affiliations:** Research Group Experimental Oto-rhino-laryngology (ExpORL), Department of Neurosciences, KU Leuven - University of Leuven, 3000 Leuven, Belgium

**Keywords:** cochlear implant, artifact rejection, neural decoding, natural speech, EEG, speech understanding

## Abstract

The speech envelope is essential for speech understanding and can be reconstructed from the electroencephalogram (EEG) recorded while listening to running speech. This so-called neural envelope tracking has been shown to relate to speech understanding in normal hearing listeners, but has barely been investigated in persons wearing cochlear implants (CI). We investigated the relation between speech understanding and neural envelope tracking in CI users.

EEG was recorded in 8 CI users while they listened to a story. Speech understanding was varied by changing the intensity of the presented speech. The speech envelope was reconstructed from the EEG using a linear decoder and then correlated with the envelope of the speech stimulus as a measure of neural envelope tracking which was compared to actual speech understanding.

This study showed that neural envelope tracking increased with increasing speech understanding in every participant. Furthermore behaviorally measured speech understanding was correlated with participant specific neural envelope tracking results indicating the potential of neural envelope tracking as an objective measure of speech understanding in CI users. This could enable objective and automatic fitting of CIs and pave the way towards closed-loop CIs that adjust continuously and automatically to individual CI users.

## 1 INTRODUCTION

Speech is characterized by fast and slow modulations. The slow modulations are also called the envelope of speech, reflecting the different syllable, word and sentence boundaries known to be essential for speech understanding (Shannon et al., 1995). Previous studies have shown that the brain tracks the speech envelope and that it is possible to reconstruct the envelope from brain responses in normal hearing listeners using electroencephalography (EEG) or magnetoencephalography (Aiken and Picton, 2008; Luo and Poeppel, 2007; Ding and Simon, 2011; Ding et al., 2015; Meyer et al., 2017). The correlation between this reconstructed envelope and the real speech envelope reflects a measure of neural envelope tracking. Recently, researchers were able to establish a link between increasing neural envelope tracking and increasing speech understanding using speech versus non-speech stimuli (Molinaro and Lizarazu, 2017), priming and vocoders (Di Liberto et al., 2018) or by adding background noise to the speech signal (Ding and Simon, 2013; Ding et al., 2014; Vanthornhout et al., 2018), underlining the application potential of neural envelope tracking as an objective measure of speech understanding.

Besides the promising results in normal hearing listeners, neural envelope tracking has been measured in listeners with a hearing impairment by Petersen et al. (2017). They showed that the amount of hearing loss could be related to neural tracking of the to-be-ignored speech, diminishing the difference between the attended and unattended speech stream in persons with increasing hearing loss.

Despite encouraging results in both normal hearing and hearing impaired listeners, for cochlear implant (CI) users, the link between neural envelope tracking and speech understanding has not been established. A CI is an implanted hearing aid that gives severely hearing-impaired to deaf persons the opportunity to (re)gain access to sound through electrical stimulation of the auditory nerve (Loizou, 1998; Wouters et al., 2015). Numerous studies have demonstrated good speech understanding for CI users in quiet, some even similar to normal hearing participants. However, when noise is added, speech understanding drops far below scores of normal hearing listeners and a large variation is seen between CI users (van Wieringen and Wouters, 2008; Shannon et al., 2011; Loizou et al., 2000). To better characterize and eventually overcome these speech understanding problems with the help of objective and automatic fitting of the CI, an objective way to measure speech understanding is of great interest. For this reason it would be useful to investigate neural envelope tracking, a potential objective measure of speech understanding, in CI users. In addition, CI users have to rely almost entirely on slow temporal information, i.e., the speech envelope, to understand speech. Therefore CI users seem excellent candidates for using neural envelope tracking.

However, a mayor problem when measuring neural envelope tracking in CI users are the stimulation artifacts in the EEG, e.g., Somers et al. (2018b); Deprez et al. (2017); Hoffman and Wouters (2010). When a CI electrically stimulates the auditory nerve, large electric potentials from the CI appear in the EEG. Just like the brain responses, the modulations in these CI artifacts follow the envelope of the presented speech signal. As a consequence it is difficult to distinguish between the desired brain response and the unwanted artifact, leading to false positives. Recently a novel artifact removal method has been developed by Somers et al. (2018b) which works by leaving out small groups of stimulation pulses during the presentation of a speech signal. This creates short artifact-free EEG windows during which the ongoing brain responses can be measured. This method was validated in CI users.

In the current study we investigated the effect of speech understanding on neural envelope tracking using EEG in CI users by applying the newly proposed artifact removal method by Somers et al. (2018b). We hypothesized that neural envelope tracking will increase with speech understanding.

## 2 MATERIAL AND METHODS

### 2.1 Participants

Eight participants aged between 46 and 75 years took part in the experiment after providing informed consent. Participants had Flemish as their mother tongue and were all experienced CI users (>8 months) using Cochlear Ltd. devices. None of them experienced tinnitus, except for S4 when the CI is off. Relevant details are shown in table 1. The study was approved by the Medical Ethics Committee UZ Leuven / Research (KU Leuven) with reference S57102.

**Table 1.** Relevant participant details.

| Name | Age (years) | Gender | CI experience | CI side | CI device | Dynamic range* | Pulse width |
| --- | --- | --- | --- | --- | --- | --- | --- |
| S1 | 58 | F | 1.1 | R | CI522 | 65 | 37 |
| S2 | 75 | M | 1.3 | L | CI522 | 61 | 37 |
| S3 | 73 | F | 0.7 | L | CI522 | 67 | 37 |
| S4 | 61 | F | 1.1 | L | CI522 | 40 | 37 |
| S5 | 58 | M | 2.0 | L | CI522 | 78 | 37 |
| S6 | 71 | M | 4.7 | L | CI24RE | 53 | 50 |
| S7 | 46 | F | 6.2 | R | CI24RE | 33 | 25 |
| S8 | 56 | F | 1.8 | R | CI512 | 62 | 37 |

\* Dynamic range is the average of all electrode specific dynamic ranges per participant.

### 2.2 Preprocessing of the stimuli

All stimuli were presented directly to the participant’s CI using a research speech processor (L34) provided by Cochlear Ltd., in combination with APEX 3 software (Francart et al., 2008) and the Nucleus Implant Communicator (NIC). To enable this direct stimulation, we converted audio files into sequences of electrical stimulation pulses using the Nucleus Matlab Toolbox (NMT) (Swanson and Mauch, 2006) in MATLAB (version R2016b) which simulates the signal processing done by a clinical CI using the ACE strategy. Such a sequence is obtained by splitting the audio waveform into frequency bands which are mapped to 22 electrodes covering a frequency range from 188 to 7938 Hz. Next, an electric pulse train is modulated with the envelope of the signal in each of these frequency bands. Finally, every modulated pulse train is sent to one of the 22 electrodes implanted inside the cochlea to stimulate the auditory nerve and enable hearing. All electrical stimulation is programmed based on the participant’s clinical settings. More concrete, the lowest audible stimulation level (threshold (T)) and the loudest stimulation level that can be comfortably listened to (C) are set per participant per electrode. Next, the amplitudes of the pulse sequences in each channel are mapped to electric currents, i.e., current units, that evoke sound sensations that are just between T and C level. Other parameter settings are pulse rate (fixed at 900 pulses per second), stimulation mode (MP1+2 = monopolar), number of active electrodes (22 electrodes, except for participant S1: 18 electrodes) and pulse width (see table 1). To vary speech understanding we varied the stimulation levels by shifting all stimulation levels between T and C level by an equal number of current units. In this way we maintain the dynamic range of the speech envelope but vary the stimulation levels and thus audibility and consequently speech understanding (figure 1).

**Figure 1.**
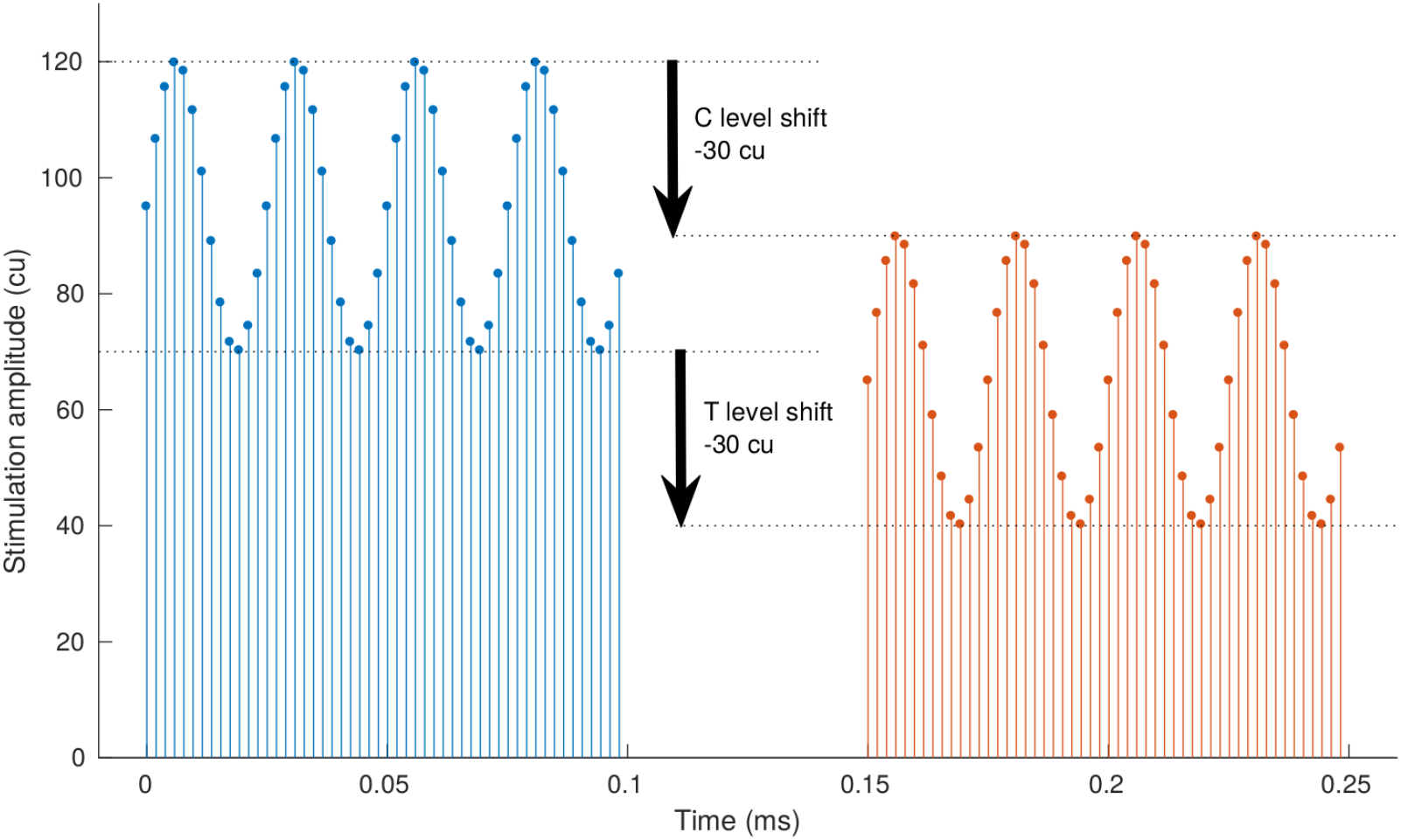
Illustration of a stimulation level shift for a sinusoidally modulated pulse train. A stimulation level shift is obtained by varying all the stimulation levels between the lowest audible (threshold (T)) and most comfortable level (C) with a fixed amount of current units (cu). Shifting the stimulation levels of the CI varies speech understanding while maintaining the dynamic range of the speech envelope.

In addition, to tackle the problem of CI stimulation artifacts when using EEG, we applied a new artifact removal method proposed and validated by Somers et al. (2018b). In brief, this method obtains EEG free of stimulation artifacts by periodically interrupting the electrical stimulation by leaving out small groups of stimulation pulses as shown in figure 2 and further referred to as a stimulus with ‘dropped pulses’. Within these interruptions artifact-free EEG can be sampled. The stimulus interruptions were 4 ms long at a rate of 40 Hz. This is short enough to preserve speech understanding and long enough to remove the artifact. Only the EEG within the stimulation gaps, i.e., artifact free EEG, is further analyzed.

**Figure 2.**
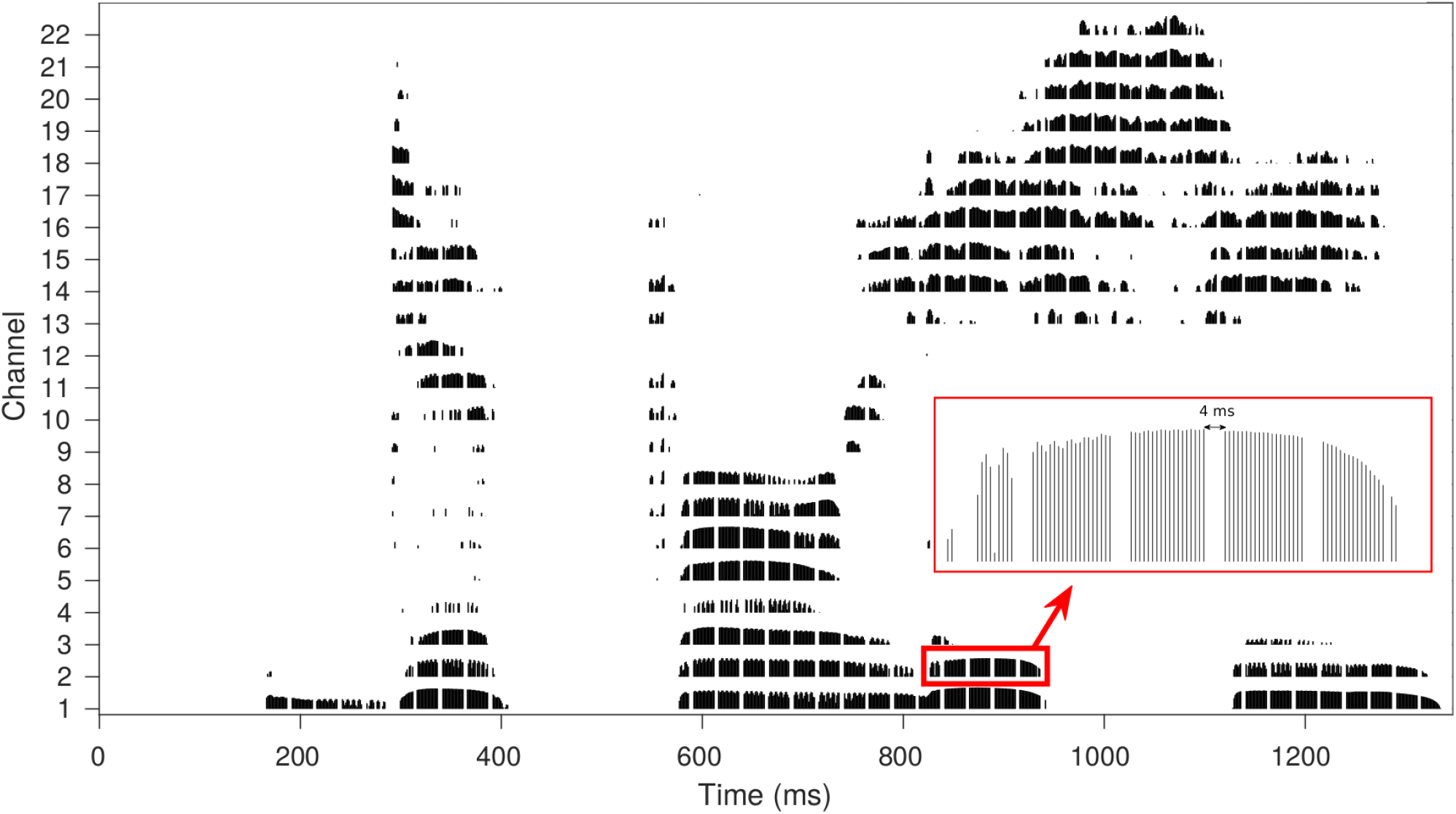
Illustration of gap insertion in the electrodogram for the Flemish word ‘politie’ (police). Every channel contains an electric pulse train modulated with the speech envelope of the corresponding frequency band. The periodically inserted gaps to obtain artifact free EEG are 4ms long at a rate of 40 Hz.

### 2.3 Behavioral experiment

Before the start of the experiment, we checked whether the participant felt comfortable listening to the stimulus with dropped pulses by presenting them with a sentence including stimulation gaps of 4 ms at a rate of 40 Hz. These values were chosen based on the findings of Somers et al. (2018b). If the participant could not tolerate the distortions induced by the gaps, we decreased the gap rate and/or length until a comfortable level was reached.

After selecting the optimal gap parameters, speech understanding was measured behaviorally in order to compare neural envelope tracking results with actual speech understanding. We used the Leuven intelligibility sentence test (LIST) (van Wieringen and Wouters, 2008), uttered in Flemish by Wivine Decoster (female speaker). This speech material is suitable to measure speech understanding in CI users or persons with a severe hearing loss due to the low speech rate (2.5 syllables/second) and the key word scoring. Every participant started with 1 training list at an input level equivalent to an acoustic signal of 60 dB SPL at the CI microphone, which will be referred to in the sequel as the baseline settings. Thereafter a number of lists, each containing 10 sentences, were presented at different stimulation level shifts to vary speech understanding. Stimulation levels were only shifted to lower values, as stimulating current units (cu) above C levels would make speech too loud and could harm the participant. Stimulation level shifts varied from 0 cu (i.e., baseline settings) to −50 cu (almost inaudible as C levels shift under original T levels for most participants depending on their dynamic range). Participants had to recall the sentence they heard. By counting the correctly recalled key words, a percentage correct per presented stimulation level shift was calculated. Additionally a list without stimulation gaps was presented at baseline settings to check the influence of inserting gaps on speech understanding.

### 2.4 EEG experiment

Recordings were made in a soundproof and electromagnetically shielded room. A 64-channel BioSemi ActiveTwo EEG recording system was used at a sample rate of 16384 Hz. Participants sat in a comfortable chair and were asked to move as little as possible during the recordings. The stories used during the EEG measurement were the stories ‘Marfoesjka en de Vorst’ (translated Russian fairytale) and ‘Luna van de boom’ (Bart Moeyaert), both narrated in Flemish by Wivine Decoster, the speaker of the LIST sentences. The stories had a total length of 48 minutes. The stories were cut in different parts: 3 longer parts of *±*8 minutes and 10 parts of *±*2.4 minutes which were presented in chronological order. The long parts were presented at baseline settings and used to train the decoder on. The short parts were presented at stimulation level shifts from a fixed list in random order, containing 0cu, −5cu −10cu, −15cu, −20cu, −30cu, −40cu, to vary speech understanding as shown in the experiment overview in figure 3. Every stimulation level shift was applied twice, on a different part of the story, to analyze test-retest reliability.

**Figure 3.**
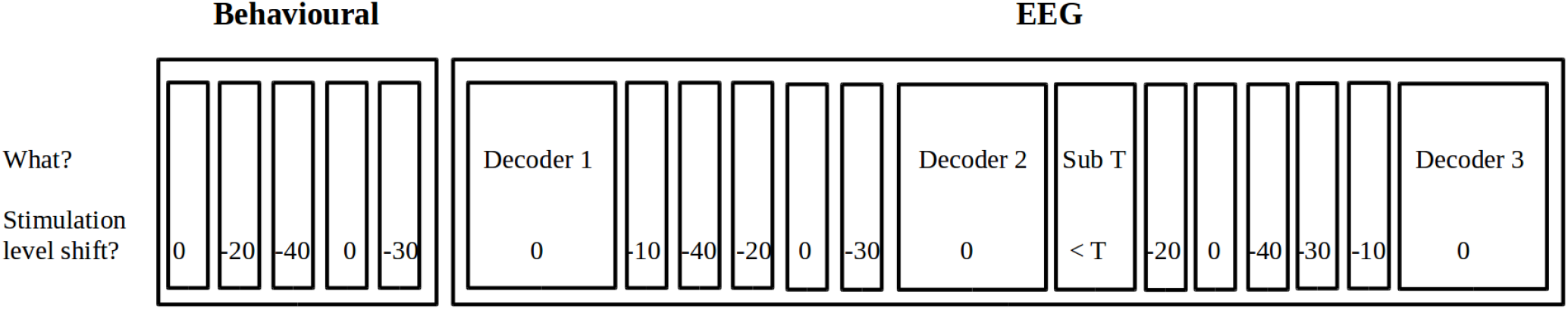
Overview of an example test session starting with the behavioral part where we measured speech understanding in a standardized way with the LIST sentences. In the second part, the EEG part, 2 stories were presented containing 3 blocks of 8 minutes without shift to train the decoder on and 10 blocks of 2.4 minutes at stimulation level shifts from a fixed list in random order, containing 0cu, −5cu −10cu, −15cu, −20cu, −30cu, −40cu, to vary speech understanding. In the middle a 5 minute part of the story was presented sub threshold (inaudible).

Additionally, we presented 5 minutes of the story below the participant’s T level (referred to as sub-T), meaning all stimulation was lower than the lowest stimulation level audible for that participant, which resulted in no sound sensation. In the sub-T condition there are CI stimulation artifacts that are highly correlated with the stimulus envelope (the CI does stimulate) in the absence of neural responses (the participant cannot hear any sound). This condition can be used to check if CI artifacts are effectively removed. If a significant neural envelope tracking response is found in this condition, this is caused by remaining CI artifacts. If no significant response is found in this condition, we can assume that the CI artifacts are removed.

To maximize the participants attention, content questions were asked after each part of the story. To measure speech intelligibility of the story, we could not ask the participant to recall every sentence similar to the behavioral LIST experiment. Therefore we used a rating method where the participants were asked to rate their speech understanding on a scale from 0 to 100% following the question ‘Which percentage of the story did you understand?’. After each trial a summary of the story presented in the previous trial was shown on the screen in front of the participant to ensure comprehension of the storyline and keep them motivated.

### 2.5 Signal processing

A measure of neural envelope tracking was calculated by correlating the stimulus envelope with the envelope reconstructed from the EEG. All signal processing was done in MATLAB (version R2016b).

#### 2.5.1 Stimulus envelope

In studies evaluating neural envelope tracking in acoustic hearing, the acoustic stimulus envelope is chosen as the reference envelope. In this study, investigating electrical hearing, using the electrical envelope as a reference is more appropriate (Somers et al., 2018b). The electrical envelope namely takes into account all the preprocessing done by the CI before the speech reaches the cochlea, making this a more realistic reference. The electric speech envelope was extracted from the stimulus by combining the magnitudes of the electrical pulses over all stimulation channels. As a next step, the electric speech envelope was band-pass filtered in the delta (0.5-4 Hz) frequency band.

#### 2.5.2 Envelope reconstruction

As a first step CI artifacts were removed by retaining only samples within the stimulus gaps as mentioned in paragraph 2.2. Other common EEG artifacts such as eye blink and muscle artifacts were removed using a multi-channel Wiener filter (Somers et al., 2018a). After artifact rejection, the signal was bandpass filtered, similar to the electric speech envelope. The mTRF Matlab toolbox (Lalor et al., 2006, 2009) was used to compute the linear decoder which reconstructs the speech envelope from the EEG recordings. As speech elicits neural responses with some delay, the linear decoder combines EEG channels and their time shifted versions to optimally reconstruct the speech envelope. If *g* is the linear decoder and *R* the shifted neural data, the reconstruction of the speech envelope *ŝ*(*t*) was obtained by *ŝ*(*t*) =∑_*n*_ ∑_*τ*_ *g*(*n, τ*)*R*(*t* + *τ, n*) with t the time ranging from 0 to *T*, *n* the recording electrode ranging from 1 to *N* and *τ* the number of post-stimulus samples used to reconstruct the envelope. The decoder was calculated using ridge regression by solving *g* = (*RR^T^*)^*−1*^(*RS^T^*) with *S* the speech envelope. As we used an integration window from 0 until 250 ms post-stimulus, the decoder matrix *g* was a 64 (EEG channels) x 11 (time delays) matrix. The decoder was created using a combination of the long segments (3 x 8 min), not including the short 2.4 minute trials.

To investigate neural envelope tracking we did two different analyses. First, to check whether we managed to eliminate the artifact from the EEG data, we did a leave-one-out cross-validation on the 24 minutes of speech at baseline settings for every sample inside the inserted stimulation gap. We hypothesized that correlations calculated for samples that contain artifacts will be higher as the artifact resembles the signal of interest. When using samples further in the gap, where the artifact has died out, correlations will be smaller, similar to correlations of previous studies with acoustic listeners, only containing the possible neural response. This leave-one-out cross-validation was done by splitting the data in equal parts of 2 minutes. The first part (2 minutes) was selected as the testing data, while the rest was concatenated to create one decoder used to reconstruct the envelope for the testing part. Next, the second part was selected as the testing data and another decoder was trained on the remaining parts et cetera. After checking if the samples were artifact-free, we concatenated the 24 minutes of EEG data at baseline settings to create one decoder per participant to apply on the 10 short segments with different stimulation level shifts, resulting in 10 outcome correlation measures per participant, i.e., 1 per trial.

### 2.6 Statistical Analysis

Statistical analysis was performed using R (version 3.3.2) software. The significance level was set at *α*=0.05 unless otherwise stated.

To analyze the behavioral results and test-retest for envelope reconstruction we used the nonparametric Wilcoxon signed-rank test for dependent samples.

To investigate the relationship between the shift in stimulation levels and neural envelope tracking or speech understanding per participant, a linear mixed effect (LME) model was constructed of neural envelope tracking/speech understanding in function of stimulation level shift (continuous variable) with a random slope and intercept per participant. To check if every chosen effect benefited the model the Akaike Information Criterion (AIC) was calculated. The model with the lowest AIC was selected and its residual plot was analyzed to assess the normality assumption of the LME residuals. Degrees of freedom (df), t-values, and p-values are reported in the results section.

To calculate the correlation between envelope reconstruction and the speech reception threshold (SRT) we used a Pearson’s correlation.

## 3 RESULTS

### 3.1 Behavioral speech intelligibility

For 7 out of 8 participants, the gap settings of 4 ms and 40 Hz were chosen. One participant (S7) reported that the speech was too distorted, a gap length of 3 ms was chosen for this participant. All participants reported that the speech with dropped pulses sounded more robotic, but speech understanding was not affected. We checked this by comparing the recall scores for LIST sentences with and without dropped pulses at baseline settings and found no significant difference between the two (p=0.7352, CI(95%) = [-13.00%; 6.00%], n=7, Wilcoxon signed-rank test). In the following analysis S8 is included to investigate the artifacts, but excluded to investigate the possible link of neural envelope tracking with speech understanding as she participated in the pilot study and only listened to the stories at baseline settings.

To investigate the effect of shifting stimulation levels on speech understanding we presented LIST sentences at different stimulation levels and asked the participants to recall the sentences. Figure 4 shows that speech understanding decreases when stimulation levels decrease for the LIST sentences (fixed effect stimulation level shift, df = 23, t= 8.74, p<0.0001, LME). During the EEG a story was presented and we asked the participants to rate their speech understanding. Similar to the LIST sentences, speech understanding for the story also decreased with decreasing stimulation levels (fixed effect stimulation level shift, df = 28, t= 5.11, p<0.0001, LME). The variation between participants was larger for the story (self-rated) than for the LIST sentences (recall) as shown in figure 4.

**Figure 4.**
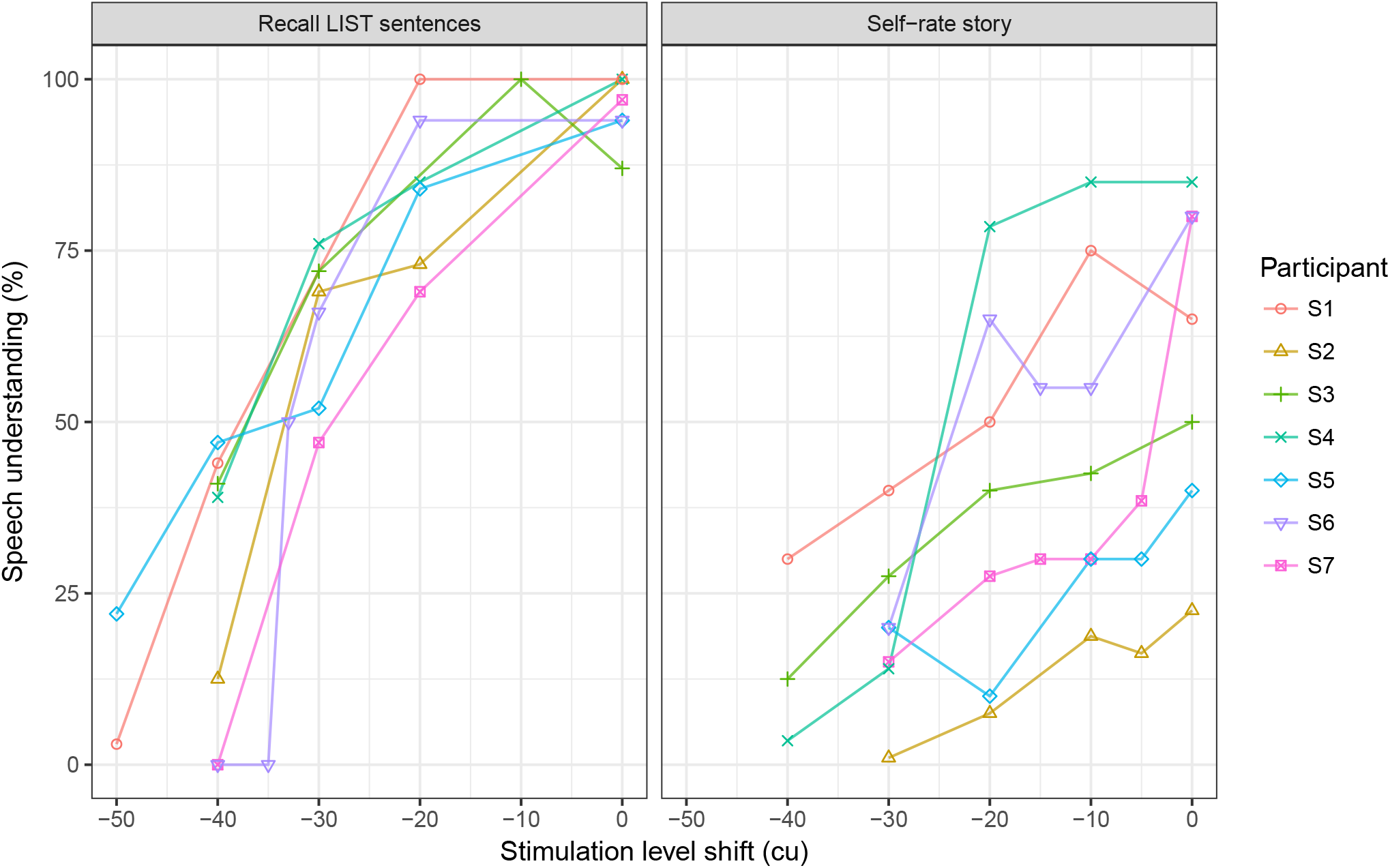
Speech understanding decreases with increasing stimulation level shift. More variation between participants is present for self-rated speech understanding of the story compared to the recalled scores of the LIST sentences.

### 3.2 Neural envelope tracking

#### 3.2.1 Influence of CI stimulation artifacts on neural envelope tracking

To investigate the presence of CI stimulation artifacts, we did two types of analysis. First we investigated the magnitude of the correlation between the real and the reconstructed envelope for each sample inside the stimulation gap. We hypothesized that correlations calculated for samples that contain artifacts will be especially high as the artifact resembles the signal of interest. When using samples where the artifact has died out, correlations will be smaller, similar to correlations of previous studies with acoustic listeners, only containing the possible neural response.

Figure 5 shows the average magnitude of the correlation for every sample inside the gap per participant at baseline settings (red color). The bold line represents the mean over participants. The correlation between the stimulus envelope and reconstructed envelope consisting of samples at the start of the gap is high. When using samples at a later point inside the gap, the correlation decreases until it reaches a plateau at the end of the gap. This asymptotic decay inside the gap is present for every participant. The circles at the right hand side of the graph indicate the average condition. This condition uses the average of the samples of the last millisecond of the gap to reconstruct the envelope, reducing the EEG noise and resulting in a more reliable reconstruction. Important to note is that one participant (S7) listened to speech with stimulation gaps of only 3 ms, resulting in deviant results in function of gap length. The results of this participant are shown on the graph as a dotted red line, but are not included to calculate the mean value of the group.

**Figure 5.**
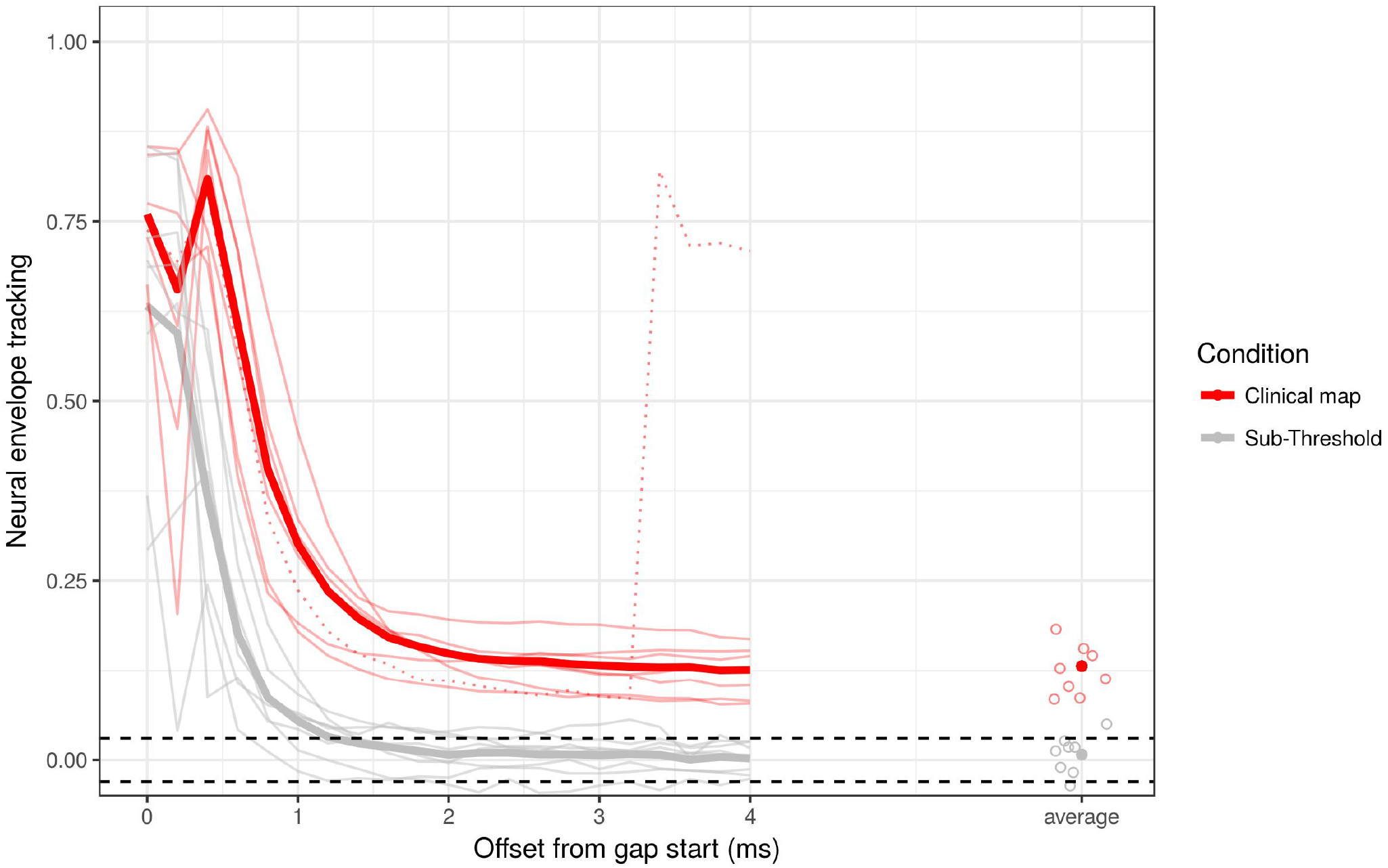
Neural envelope tracking results over the gap length. The red lines are the mean result per participant at baseline settings. The black lines are the result of the condition where speech was presented sub threshold. The bold lines represent the mean over participants. The dashed lines are the 95% significance level of the correlation. The circles at the right hand side show the results of the average condition. The dotted red line shows the results of S7 with a deviant gap length.

Next we calculated the correlation between the real and reconstructed envelope in the sub-T condition. In this condition the CI was stimulated, but the recipient could not hear any sound. Results of this analysis are shown in figure 5 in grey. Correlations at the beginning of the gap are still significant, but when taking a sample far enough from the start, the correlation is below or within significance level for 6 out of 7 participants. The significance level of the correlation is indicated by the dashed horizontal lines. It is calculated by correlating random permutations of the real and reconstructed envelope 1000 times and taking percentile 2.5 and 97.5 to obtain a 95% confidence interval.

#### 3.2.2 Relation between speech understanding and neural envelope tracking

To investigate whether speech understanding is related with neural envelope tracking in CI users, we varied the stimulation levels by reducing them with a fixed value as shown in figure 1. To measure neural envelope tracking, we calculated the Spearman correlation between the reconstructed envelope and the stimulus envelope for an average of the samples within the last millisecond of the presented stimulation gap to exclude artifacts. Conducting a test-retest analysis showed no significant difference between test and retest correlations (p=0.59, CI(95%) = [-0.011; 0.019], Wilcoxon signed-rank test), therefore we averaged the correlation of the test and retest conditions resulting in one correlation per participant per stimulation level shift, except for participant S5 who only participated in the test condition.

Figure 6 shows that the more the stimulation levels (lower x-axis) decrease, the more the correlation between the real and the reconstructed envelope, i.e., neural envelope tracking, also decreases (fixed effect stimulation level shift, df = 28, t= 4.60, p=0.0001, LME). In addition, the results of the sub-T condition are also shown. Similar to the sub-T results in figure 5, neural envelope tracking in the sub-T condition is below or within significance level for 6 out of 7 participants. As an extra factor average speech understanding across participants of the LIST sentences is also included in the figure on the upper x-axis to show the interplay between decreasing stimulation levels, decreasing speech understanding and decreasing neural envelope tracking (fixed effect speech understanding on neural envelope tracking, df = 28, t= 3.56, p=0.0013, LME).

**Figure 6.**
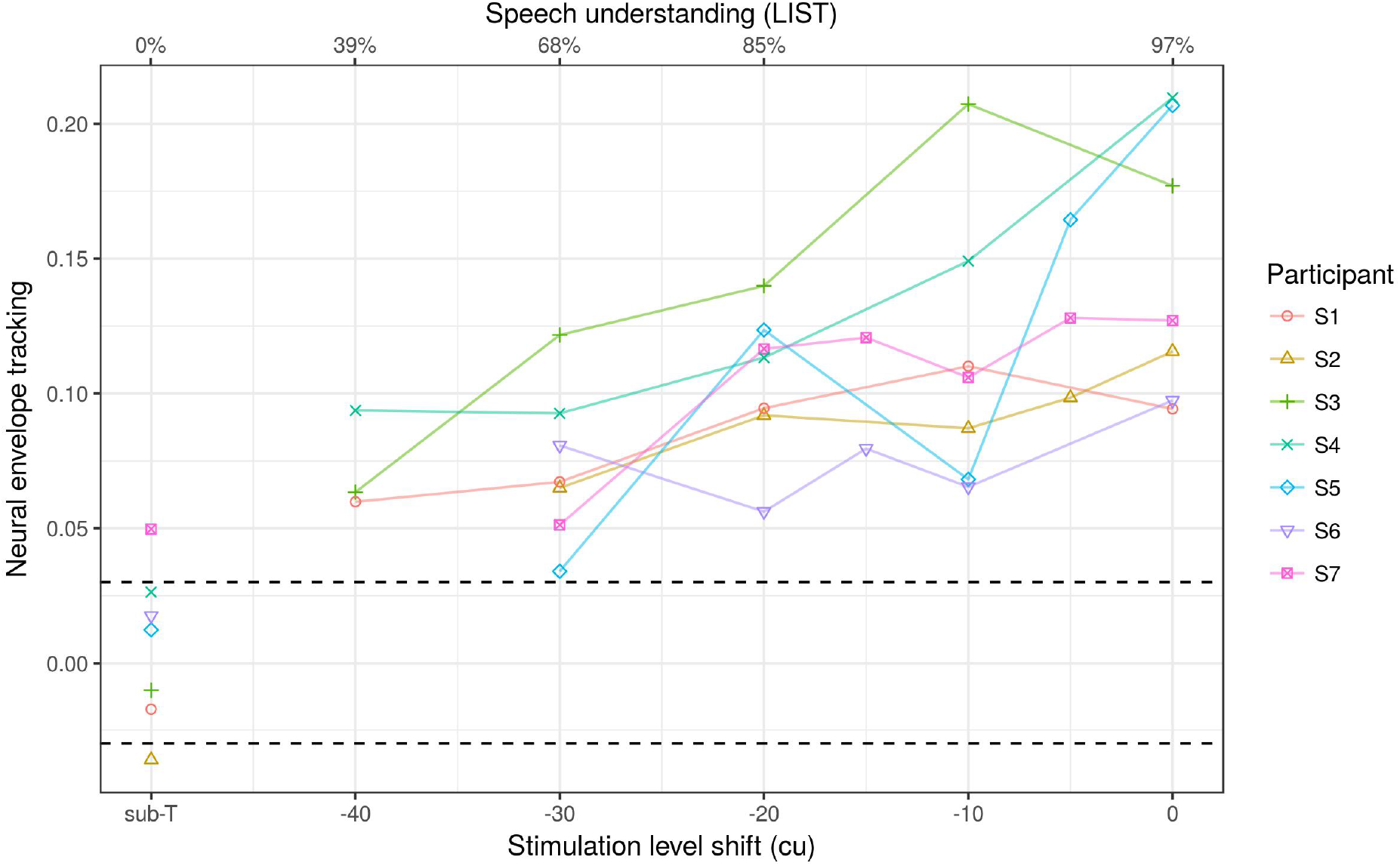
Neural envelope tracking increases with increasing stimulation level shift. The dashed black lines are the 95% significance level of the correlation. The values for speech understanding are the median values for the LIST sentences over participants.

In the previous analysis we investigated the relation between speech understanding and neural envelope tracking on a group level. To have a closer look at how participant specific speech understanding relates to neural envelope tracking, we calculated the speech reception threshold (SRT) per participant, i.e., the stimulation level shift yielding 50% speech understanding. This 50% point was calculated by fitting a linear function on the data and solving the equation: 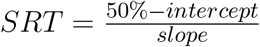 as shown in figure 7A. Next, we correlated the SRT of every participant with the neural envelope tracking score on the 24 minutes of speech at baseline settings, represented as circles on the right hand side in figure 5. The more negative the SRT, the better the participant understands speech, the higher neural envelope tracking at baseline settings (Pearson correlation = −0.76, p=0.048, figure 7B).

**Figure 7.**
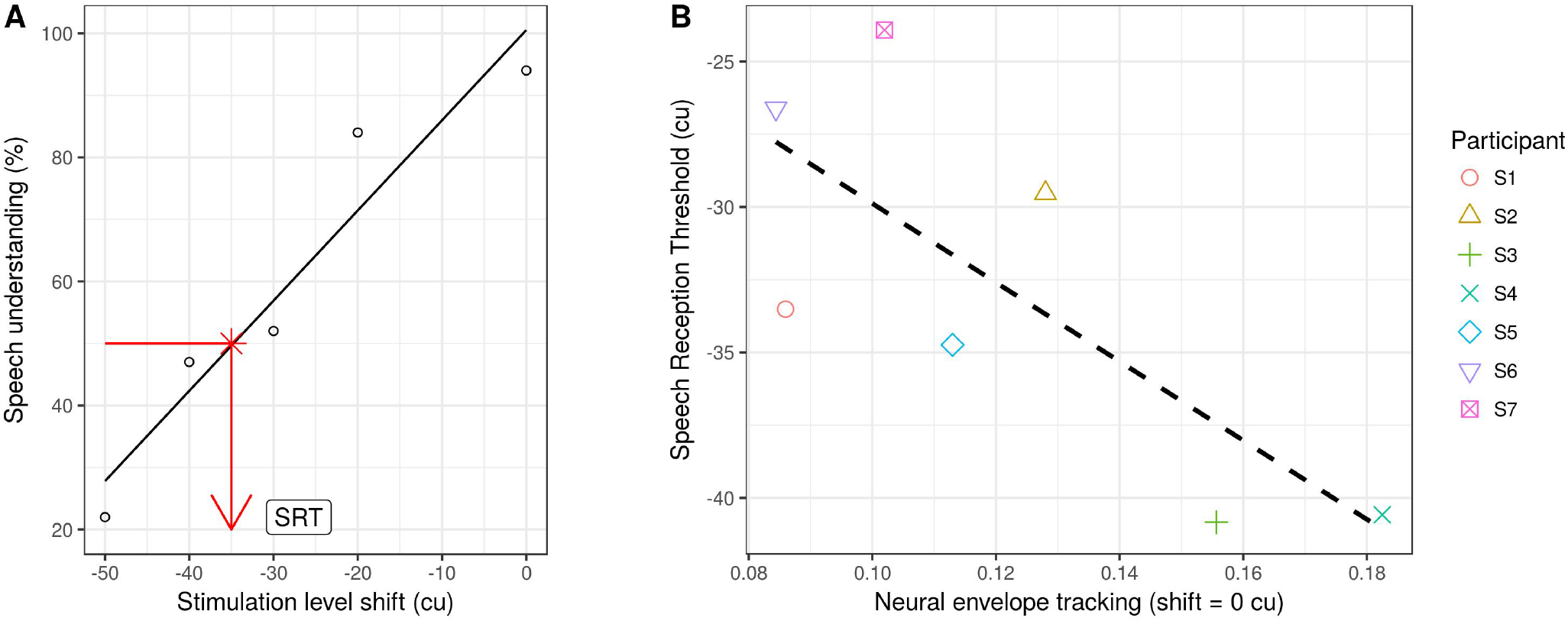
Neural envelope tracking correlates with speech understanding. Panel A shows how the behavioral speech understanding results of 1 participant (S5). A linear function is fitted through the data and the stimulation level shift corresponding to 50% speech understanding is labeled as the SRT. Panel B shows that the better (negative) the SRT, the higher neural envelope tracking is at baseline settings.

## 4 DISCUSSION

In this study we investigated if neural envelope tracking is related to speech understanding in CI users similar to normal hearing listeners. To that end, we recorded the EEG of 8 CI users listening to a story at varying levels of speech understanding by shifting the stimulation levels. An envelope reconstruction analysis was conducted and compared to speech understanding results. We found increasing neural envelope tracking with increasing stimulation levels and corresponding speech understanding which supports the hypothesis that neural envelope tracking is related with speech understanding in CI users.

### 4.1 Speech understanding influenced by stimulation level shifts

As a first step we checked if the chosen method to vary speech understanding, i.e., shifting stimulation levels, achieved the desired outcome by presenting several lists of LIST sentences at various stimulation level shifts. Similar to results in normal hearing listeners where decreasing stimulus intensity is accompanied by decreasing speech understanding (van Wieringen and Wouters, 2008), we found that decreasing stimulation levels resulted in decreasing speech understanding. In addition, speech understanding did decrease gradually with decreasing stimulation levels, giving the opportunity to investigate neural envelope tracking at a wide range of speech understanding levels. Next we checked if the introduced stimulation gaps, necessary to remove the artifact, affected speech understanding. This was not the case, however they did affect the quality of the speech signal, possibly resulting in more listening effort which has to be taken into account when comparing results of CI users to results of normal hearing listeners.

Besides the standardized recall test of the LIST sentences, we also asked the participants to rate their speech understanding of the presented story during the EEG. As shown in figure 4, the variation between participants is larger for the self-rated story than for the recalled LIST sentences. This can be explained by the different way speech understanding was measured with more reliable results for the recalled LIST sentences, indicating the importance of standardized speech tests in addition to self-rated measures.

### 4.2 Influence of CI stimulation artifacts on neural envelop tracking

To remove CI stimulation artifacts we used a validated artifact removal technique by Somers et al. (2018b) which leaves out small groups of stimulation pulses during the presentation of a speech signal. We were able to show that the correlation inside the gap between the stimulus envelope and reconstructed envelope decreased when using samples further in the gap until it reached a plateau at the end of the gap (figure 5). Although it is likely that the artifact was removed because of the present decay over the gap length, this can not be guaranteed. Despite this clear decay in all participants, one participant (S7) showed significant, but very low, neural envelope tracking in the sub-T condition until the end of the gap where no neural envelope tracking was expected. This could mean that a gap of 4 ms in this participant was not enough for the stimulus artifact to decay. Another explanation could be that although the stimulation was sub-T, the participant still perceived some sounds, resulting in very small neural responses.

### 4.3 Relation between speech understanding and neural envelope tracking

Next, we investigated the relation between neural envelope tracking and speech understanding by varying the stimulation levels of the CI. Similar to research in normal hearing listeners that showed increasing neural envelope tracking with increasing speech understanding (Molinaro and Lizarazu, 2017; Ding and Simon, 2013; Ding et al., 2014; Vanthornhout et al., 2018; Di Liberto et al., 2018), we showed similar results in CI users although using a different approach. We varied the stimulation levels of the CI because using speech versus non-speech stimuli (Molinaro and Lizarazu, 2017) or priming (Di Liberto et al., 2018) would only result in 2 speech understanding levels. Furthermore, adding background noise to the speech signal (Ding and Simon, 2013; Ding et al., 2014; Vanthornhout et al., 2018) would not be compatible with the artifact removal method as the noise would constantly be interrupted. Therefore we decided to directly manipulate a parameter of the CI, namely the stimulation levels. By doing so, we not only varied speech understanding but also investigated the effect of adjusting a CI parameter based on neural envelope tracking which demonstrates the feasibility of fitting a CI based on this measure. Besides the group analysis, we also investigated the relation between speech understanding and neural envelope tracking on a participant specific level similar to Vanthornhout et al. (2018). We were able to show that participants with good speech understanding (good SRTs) had enhanced neural envelope tracking at baseline settings, again showing the potential of neural envelope tracking as an objective measure of speech understanding ion CI users.

A potential confound in our study is loudness. Varying stimulation levels not only varies speech understanding, it additionally affects the loudness of the stimulation. Therefore, it is difficult to distinguish if neural envelope tracking decreased with decreasing stimulation levels because of speech understanding or loudness. However Ding and Simon (2012) showed no difference in neural envelope tracking for a variation in stimulus intensity over 16 dB. Nevertheless, even if the found effect would be influenced by loudness in addition to speech understanding, this would still be an interesting result in the context of objective fitting, showing that neural envelope tracking can be used to adjust the parameter settings of the CI.

### 4.4 Implications for applied research

This study is the first to show a link between neural envelope tracking and speech understanding in CI users, indicating the potential of neural envelope tracking as a measure of speech understanding in CI users. Further research in this field could enable the development of an objective measure of speech understanding in CI users with application potential in the field of objective clinical measures and neuro-steered hearing aids.

## 5 CONCLUSION

This study confirms that neural envelope tracking responses can be found in CI users in response to running speech using appropriate CI artifact removal methods. Furthermore, these responses become weaker as the stimulus is presented at less intelligible stimulation levels. Neural envelope tracking can serve as a measure of speech understanding that directly relates to settings of the CI, and thus has application potential in objective and automatic fitting of CIs.

## Acknowledgements

The authors would also like to thank Wivine Decoster for narrating the stories.

